# Revisiting the decay of scientific email addresses

**DOI:** 10.1101/633255

**Authors:** Raul Rodriguez-Esteban, Dina Vishnyakova, Fabio Rinaldi

**Affiliations:** Roche Pharmaceutical Research and Early Development, Roche Innovation Center Basel, Grenzacherstrasse 124, 4070 Basel, Switzerland; University of Zurich, Institute of Computational Linguistics and Swiss Institute of Bioinformatics, Andreasstrasse 15, Zürich, CH-8050; Dalle Molle Institute for Artificial Intelligence Research, Galleria 2, Via Cantonale 2c, CH-6928 Manno (Lugano), Switzerland

## Abstract

Email is the primary means of communication for scientists. However, scientific authors change email address over time. Using a new method, we have calculated that approximately 18% of all authors’ contact email addresses in MEDLINE are invalid. While an unfortunate number, it is, however, lower than previously estimated. To mitigate this problem, institutions should provide email forwarding and scientific authors should use more stable email addresses. In fact, a steadily growing share already use free private email addresses: 32% of all new addresses in MEDLINE in 2018 were of this kind.

## Introduction

It is well known to online marketers that email addresses go “stale” (i.e., become invalid) over time. This is a phenomenon that also affects corresponding email addresses from scientific articles. Wren et al. (2006) estimated that 24% of all contact email addresses in MEDLINE become invalid within a year of publication and that that percentage approaches 50% within 5 years. In the context of a research project about author disambiguation (Vishnyakova et al., 2019), we revisited this topic using a different approach to that taken by Wren et al. (2006).

## Results

We emailed 265 authors randomly selected from the MEDLINE database. Out of those emails, we received 52 (20%) bounce notices (Fig. 1). As one would expect, the bounce rate increased with the age of the article in which the email appeared. Modeling the bounce statistics with a time-dependent Bernouilli process (see Methods) we calculated that roughly 2.1% of all contact emails in MEDLINE become invalid every year. Since there are 3,283,151 unique contact email addresses in the MEDLINE database (Nov. 19th, 2018), this means that roughly 69,000 of those addresses will go stale within a year. Using our model we also estimated that 18% of all emails in MEDLINE are currently invalid. This number will grow quickly because more than half of all email addresses in MEDLINE have only been added since 2011.

**Figure 1.**
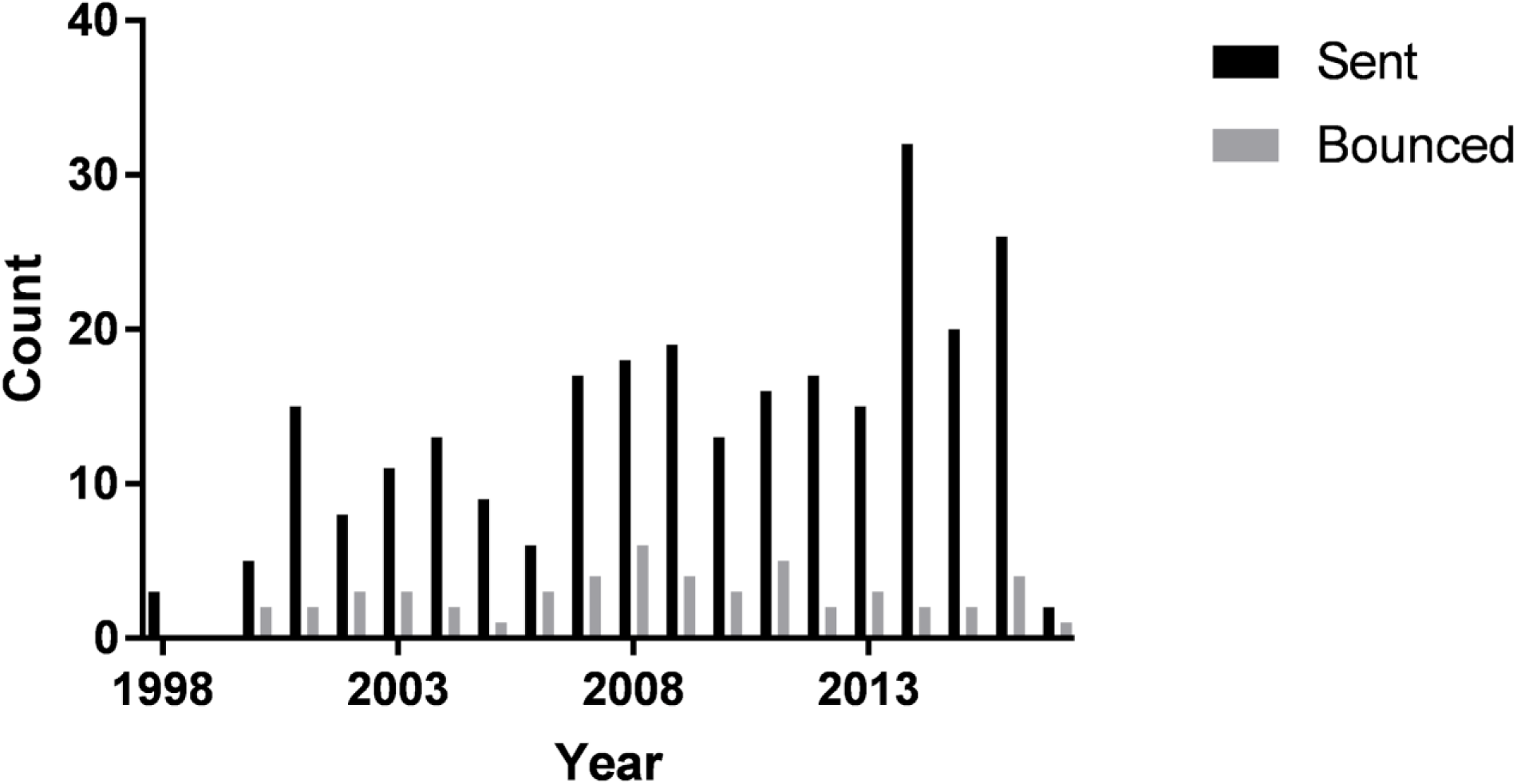
Bounce statistics. Number of emails sent and number bounced vs. year when the email address first appeared in MEDLINE (using the MEDLINE baseline 2016 as reference).

We also investigated the use of email providers in MEDLINE and noted that the share belonging to free providers has been growing, as already noticed by Kozak et al. (2015). In fact, 32% of all new email addresses in 2018 were from free email providers (Fig. 2).

**Figure 2.**
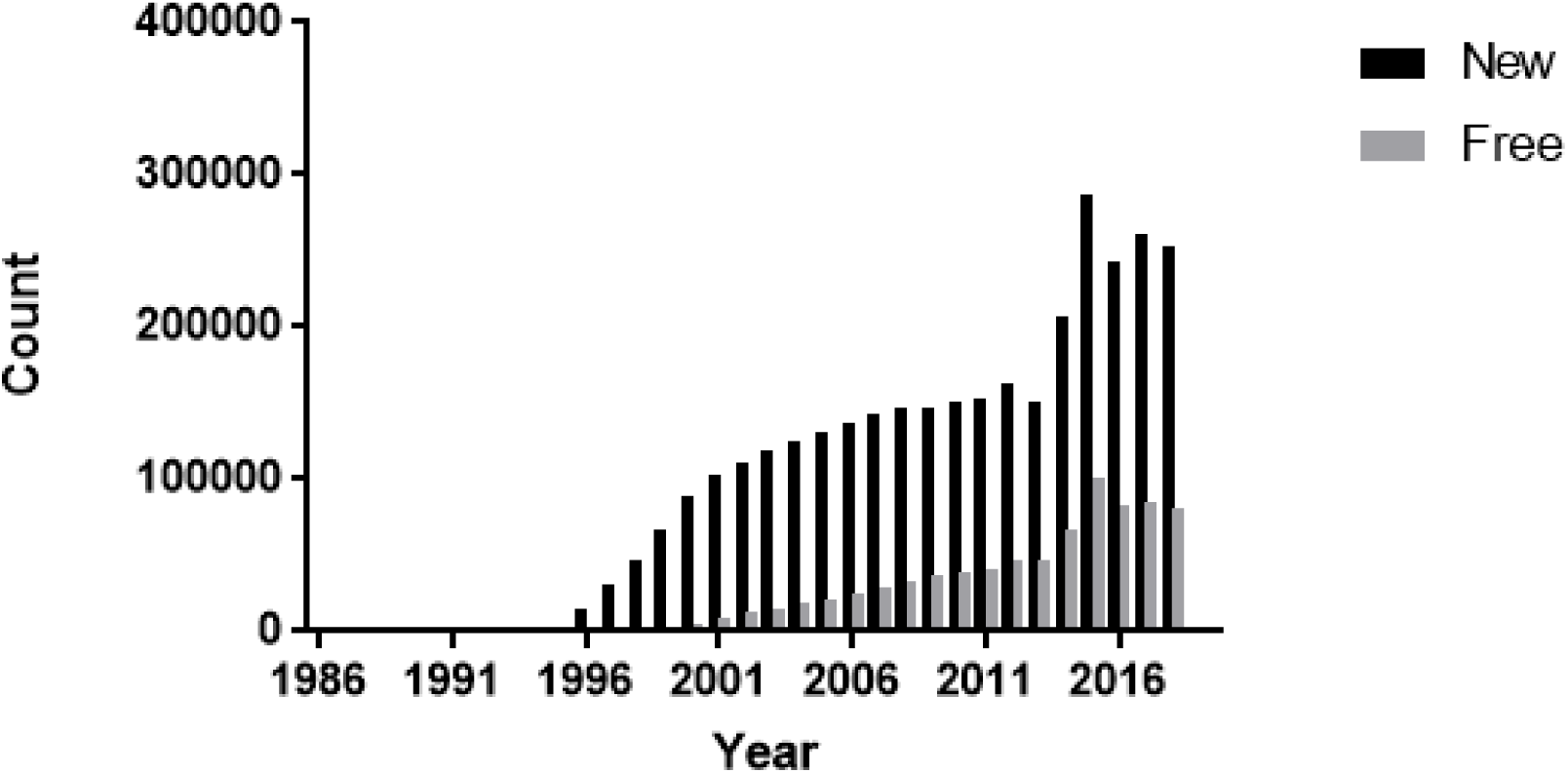
Email address statistics in MEDLINE as of Nov. 2018. The figure depicts new email addresses appearing in the MEDLINE database per year. “Free” email addresses correspond to free email providers.

## Discussion & Conclusion

Our results present a less bleak picture than that depicted by Wren et al. (2006). That study was based on responses only from email servers providing meaningful information to automatic queries. Email servers, however, have become less and less responsive over time as a protection against spammers. We believe that bounce notices in response to manually-sent emails are more reliable to gauge email address validity, particularly if the responding email server does not deem the incoming email suspicious of being spam. The method used for this study requires, moreover, less time and computation than the one used in Wren et al. (2006).

Contact email addresses from scientific articles go stale for various reasons. A common problem is the use of professional email addresses that do not get redirected after a change of employer--particularly in the corporate world. A solution for this would be for institutions to provide email redirection for all its departing scientists. Another way to tackle this problem would be for authors to use more portable email addresses, such as those offered by free email providers. If journals encouraged authors to use email addresses that are independent of the author’s current employer, this could help remove any potential bias against the use of non-institutional addresses. Yet another solution would be to enable existing online scientific directories, such as ORCID, to allow direct contact with scientists. Unfortunately, such directories are not yet in widespread use.

The ability to contact scientific authors is essential for a scientist’s daily work in tasks such as reproducing published work or exchanging reagents and materials. Readers who try to contact invalid email addresses are left to scramble for up-to-date contact information. Thus, scientific authors should be more mindful of the contact information they provide to enable the process of science to move forward and perhaps increase their scientific impact (Cokol et al., 2007; Cokol and Rodriguez-Esteban, 2008). In the age of electronic communication and online presence management there are widespread technological solutions to improve the connection between the readers and writers of scientific publications.

## Methods

Personalized emails were sent individually from a Gmail corporate account to scientific authors in November 2018. The email addresses contacted were randomly selected from the MEDLINE baseline 2016, which includes publications from 2017. The addresses were extracted from the Affiliation field from each MEDLINE record. We did not extract emails from the Abstract/Contact section or other sections, which only appear in some MEDLINE records. (We estimate that emails outside of the Affiliation section represent less than 1% of all emails in MEDLINE.) The reference date for each record was the publication date (PubDate).

Overall email address statistics corresponded to MEDLINE records up to Nov. 19th, 2018. Free email providers were taken from a list of 4,316 domains maintained in https://github.com/willwhite/freemail/blob/master/data/free.txt. Note that MEDLINE includes records with future publication dates, therefore statistics for 2018 include articles to be published in 2019.

The probability of an email address becoming invalid was derived with a linearly time-dependent Bernoulli process in which the probability of *k* emails bouncing out of *n* emails sent is defined by the equation

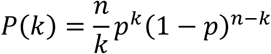

in which *p* is a linear function of time,

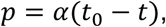

where *t*_*0*_ corresponds to November 2018 and α is a linear coefficient.

Scripts used for this analysis are available at: https://github.com/raroes/stale-emails

